# Super resolution imaging of a distinct chromatin loop in human lymphoblastoid cells

**DOI:** 10.1101/621920

**Authors:** Jacqueline Jufen Zhu, Zofia Parteka, Byoungkoo Lee, Przemyslaw Szalaj, Ping Wang, Karolina Jodkowska, Jesse Aaron, Teng-Leong Chew, Dariusz Plewczynski, Yijun Ruan

**Affiliations:** The Jackson Laboratory for Genomic Medicine, 10 Discovery Drive, Farmington, CT 06030, USA; Department of Genetics and Genome Sciences, University of Connecticut Health Center, 400 Farmington Avenue, Farmington, CT 06030, USA; Centre of New Technologies, University of Warsaw, S. Banacha 2c, 02-097 Warsaw, Poland; Faculty of Mathematics and Information Science, Warsaw University of Technology, Warsaw, Poland; Center for Bioinformatics and Data Analysis, Medical University of Bialystok, Jana Kilinskiego 1, 15-089 Bialystok, Poland; I-BioStat, Hasselt University, Agoralaan D, 3590 Diepenbeek, Belgium; Advanced Imaging Center, Janelia Research Campus, Howard Hughes Medical Institute, 19700 Helix Drive, Ashburn, VA 20147, USA; National key laboratory of crop genetic improvement, College of Life Sciences and Technology, Huazhong Agricultural University, Wuhan, Hubei 430070, China

## Abstract

The three-dimensional genome structure plays a fundamental role in gene regulation and cellular functions. Recent studies in genomics based on sequencing technologies inferred the very basic functional chromatin folding structures of the genome known as chromatin loops, the long-range chromatin interactions that are often mediated by protein factors. To visualize the looping structure of chromatin we applied super-resolution microscopy iPALM to image a specific chromatin loop in GM12878 cells. Totally, we have generated six images of the target chromatin region at the single molecule resolution. To infer the chromatin structures from the captured images, we modeled them as looping conformations using different computational algorithms and then evaluated the models by comparing with Hi-C data to examine the concordance. The results showed a good correlation between the imaging data and sequencing data, suggesting the visualization of higher-order chromatin structures for the very short genomic segments can be realized by microscopic imaging.

## Introduction

How chromatin is organized in cell nucleus is a historically mysterious question. It is known that DNA is packed in different levels to allow meters-long linear DNA to be condensed in micrometers-sized nucleus. Bound by histone proteins, 146 base pairs (bp) of DNA form nucleosomes (***Luger et al., 1997; Tsunaka et al., 2005***) that are connected by dozens of bp of linker DNA, appearing as a “beads on a string” structure (***Olins and Olins, 1974; Kornberg, 1974; Oudet et al., 1975; Finch and Klug, 1976; Bustin et al., 1976; Leuba et al., 1994***). The 10 nm “beads on a string” DNA fiber is then folded into higher-order chromatin structures for further chromatin compaction. However, the organization of higher-order chromatin structures was elusive for tens of years. Although the 30 nm chromatin fiber was observed and suggested to be the next organizational level of the 10 nm fiber, it is now debatable whether it exists *in vivo* (***Felsenfeld and Groudine, 2003; van Holde and Zlatanova, 2007; Nishino et al., 2012***). Recently, technologies combining biochemistry and high-throughput sequencing such as Hi-C (***Lieberman-Aiden et al., 2009***) and ChIA-PET (***Fullwood et al., 2009***) have been developed to characterize genome-wide landscape of long-range chromatin interactions (usually from several kilobases (kb) to hundreds of kilobases) that are considered as the basis of higher-order chromatin organization. Chromatin interactions suggest the looping structure of chromatin, describing DNA loci that are in close spatial proximity even though they are located far away in linear genomic distance. Based on chromatin interactions, more complex megabase-sized structures such as topologically associating domains (TADs) (***Dixon et al., 2012; Ricci et al., 2015; Maeshima et al., 2014***) and CTCF-mediated chromatin contact domains (CCDs) (***Tang et al., 2015***) were predicted. Importantly, these high-order structures were shown to be involved in transcription regulation (***Pope et al., 2014; Apostolou and Thanos, 2008; Ling et al., 2006***) and disease development (***Lupiáñez et al., 2015, 2016***), suggesting critical biological importance. However, the visualization of these hypothesized higher-order chromatin structures *in situ* remains difficult mainly due to the resolution limitation of conventional microscopy. While electron microscopy has been previously used to visualize the ultrastructures and 3D organization of chromatin directly (***Mahamid et al., 2016; Ou et al., 2017***), it lacks specificity to investigate transcription associated chromatin structures. Fortunately, super-resolution light microscopy has made it possible to achieve this goal. Recently, stochastic optical reconstruction microscopy (STORM) has been used to image chromatin folding of TAD and different epigenetic states inferred by Hi-C (***Wang et al., 2016; Boettiger et al., 2016***). Given these developments, there comes now a more intriguing challenge to visualize a distinct chromatin loop. In this study, we applied interferometric fluorescent super-resolution microscopy (iPALM) combined with DNA fluorescence in situ hybridization (FISH) to visualize a distinct chromatin loop occurring frequently in human lymphoblastoid cells inferred by Hi-C and ChIA-PET chromatin contact data, which enabled us to resolve the very fine chromatin looping structures within 33kb of DNA.

## Results

### The chromatin landscape and DNA FISH design of the target loop region

Hi-C and ChIA-PET contact data showed chromatin loops at various genomic loci and length scales. As a target chromatin region we selected a 13kb long, high frequent ***Table 1*** loop mediated by both CTCF and RNAPII in GM12878 cells (***Figure 1***b). The loop is located at the T-cell receptor alpha (TCRA) locus on Chromosome 14, where V(D)J recombination takes place during T-lymphocyte development and has been studied in mouse T cells (***Seitan et al., 2012, 2013; Shih et al., 2012***). A CTCF-and cohesin-binding site is located between the TCRA locus and the neighbouring housekeeping gene DAD1. It is already suggested that this site functions as an insulator, as the depletion of cohesin leads to increased transcription of DAD1 (***Seitan et al., 2011***). Interaction between TCRA enhancer and Dad1 was shown to occur in both mouse T-and B-lymphocytes, although in the latter it was slightly weaker (***Shih et al., 2012; Seitan et al., 2011***). The conservation of TCRA locus in mouse and human (***Glusman et al., 2001***) implies a similar chromatin conformation in human lymphocytes with mouse. As expected, in our GM12878 ChIA-PET data, one anchor of the loop overlaps with TCRA enhancer, and the second one is situated in Dad1 gene body, which is similar to the observation in mouse T-cells, suggesting the insulation function of the target loop in human cells, adding biological meaning to the study on its detailed structure.

**Table 1.** GM12878 CTCF ChIA-PET top five strongest loops. The iPALM target loop, highlighted by bold font, is the third strongest CTCF loop over the whole genome. The total CTCF loop number is 42297, and the average PET count is 32.8. The minimum PET count of loops is 4.

| Chromosome | Start | End | Chromosome | Start | End | PET |
| --- | --- | --- | --- | --- | --- | --- |
| chr15 | 22436211 | 22436212 | chr15 | 22461064 | 22461065 | 5523 |
| chrX | 9963956 | 9963957 | chrX | 10087576 | 10087577 | 5239 |
| **chr14** | **23026053** | **23026054** | **23039387** | **23039388** | **chr14** | **2726** |
| chr11 | 130305008 | 130305009 | chr11 | 130732043 | 130732044 | 1644 |
| chr12 | 57607189 | 57607190 | chr12 | 57633076 | 57633077 | 1409 |

**Figure 1.**
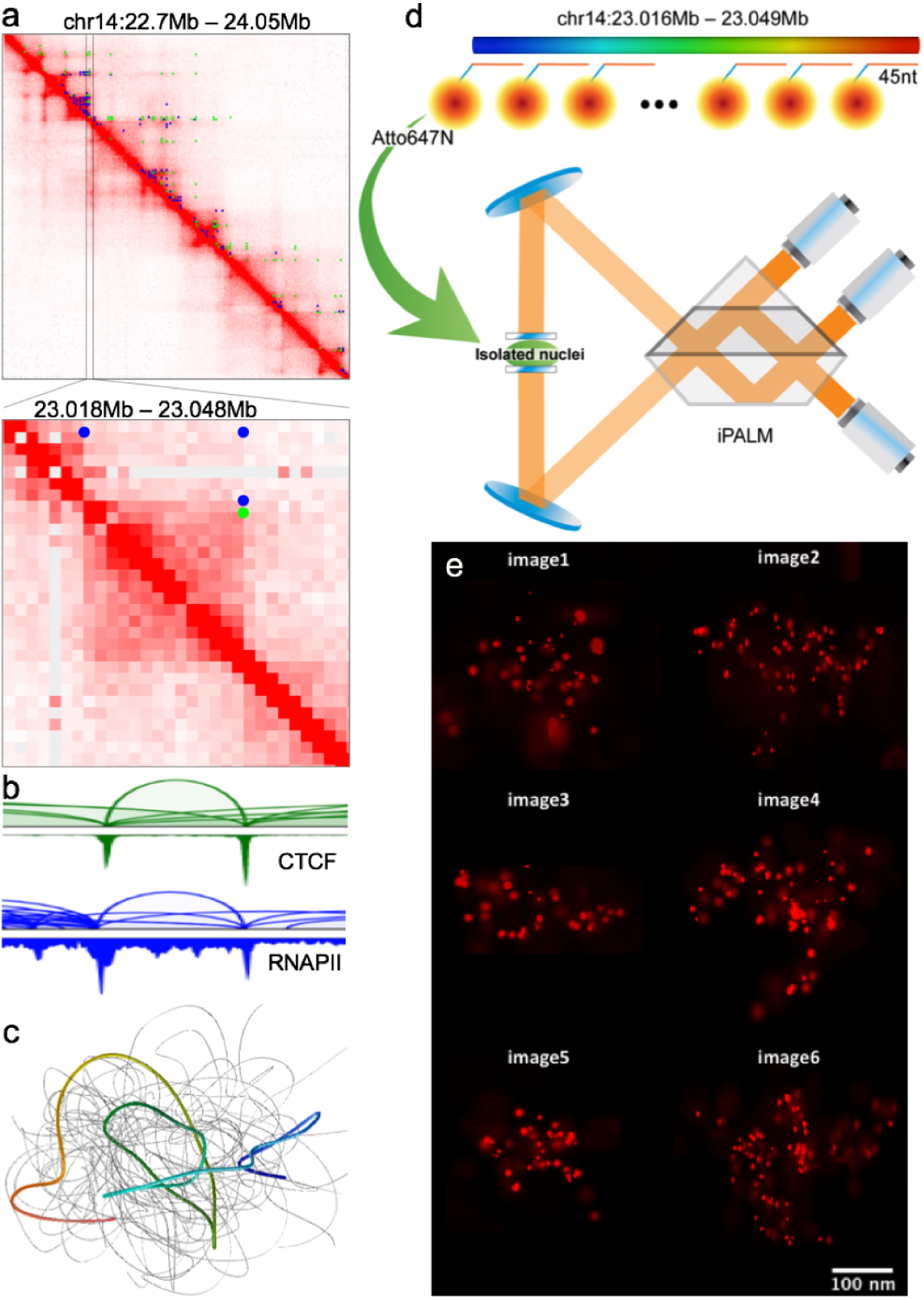
iPALM method to visualize a distinct chromatin loop. a. GM12878 Hi-C contact map (***Rao et al., 2014***) for chromosome 14: 22.7 – 24.05Mb (5kb resolution with balanced normalization, top) and zoomed-in target loop region (Chr14: 23.018 – 23.048Mb, 1kb resolution with balanced normalization, bottom). ChIA-PET loops (green for CTCF and blue for RNAPII) are also presented on top of the upper diagonal area of the contact map. b. GM12878 ChIA-PET genome browser view (***Tang et al., 2015***) for the target loop region (Chr14: 23.018 – 23.048Mb). CTCF loops and peaks (green) and RNAPII loops and peaks (blue) are presented. c. 3D chromatin models for the target loop region using Hi-C with multidimensional scaling method (***Szalaj et al., 2016; Szalaj et al., 2016***). Ensemble of 100 structures is presented. One typical model with a visible a loop structure is highlighted (rainbow color) d. Schematics of the iPALM method. The probe set contains 336 DNA oligos tagged with Atto647N designed to stain across the target loop region (Chr14: 23016081 – 23048740). e. Six observed iPALM images. **Figure 1–video 1**. Three-dimensional projection of image 1. **Figure 1–video 2**. Three-dimensional projection of image 2. **Figure 1–video 3**. Three-dimensional projection of image 3 **Figure 1–video 4**. Three-dimensional projection of image 4 **Figure 1–video 5**. Three-dimensional projection of image 5 **Figure 1–video 6**. Three-dimensional projection of image 6

To keep the integrity of the loop in consideration of staining efficiency, we extended the selected region by 10kb on both sides. The resulting 33kb chromatin region is presented in both Hi-C heat map (***Figure 1***a) and ChIA-PET browser view (***Figure 1***b). The surrounding epigenomic landscape of this region is shown in ***Figure 1–Figure Supplement 1***. We modeled the conformation of the target region using sequencing data (***Figure 1***c). To this end, we used multidimensional scaling to estimate 3D chromatin structure of the loop region from Hi-C data. At the same time we applied iPALM to visualize it (***Figure 1***d). Oligopaints probe (***Beliveau et al., 2012***) for DNA FISH was designed to target on the chromatin (***Figure 1***d) specifically by avoiding DNA repeats, resulting a staining density of about 10 oligos/kb, allowing us to visualize the target chromatin as a dot by conventional microscope, though lacking details of the structuring, facilitating target localizing with iPALM imaging.

### Fine structures of the target chromatin revealed by iPALM imaging

We applied iPALM (***Shtengel et al., 2009***) to image samples stained with Atto647N tagged probes (MYcroarray), which achieved single molecule resolution (***Figure 1***e). We acquired six high quality images for the target chromatin region (***Figure 1***e).

Briefly, samples were imaged at 30-50ms exposure, under 3kW/cm^2^ of 647nm laser excitation and 100W/cm^2^ 405nm laser activation for 25,000 frames to capture blinking Atto647N molecules. Data was imported into the PeakSelector software package (Janelia Research Campus), which registers three camera images with respect to each other, calibrates the intensity across each camera as a function of z-position, and localizes each blinking molecule in 3D. Further, fiducial nanoparticles embedded in the coverslip allow for drift correction after acquisition and localization to maximize image resolution. After the processing, spatial positions were filtered based on localization uncertainty in all three dimensions and data were rendered as 3D TIFF stacks for further analysis. Each image (***Figure 1***e) shows one copy of the 33 kb chromatin target. The well-separated red dots represent single fluorescent molecules bound along the chromatin. From the images, we can infer that the dots are not randomly distributed but ordered in some way to form featured spatial conformation. We then characterized each image by dot count, volume of the image, and the minimal distance between dots (***Figure 2***a, b, c, ***Figure 2–Figure Supplement 1***). Volume of the image was estimated by calculating the volume of convex hull for dots identified from iPALM images (***Jones et al., 2014***). On average, there are 68 dots per image. The minimal distance between dots is in the range of 0-60nm, and the average image volume is 0.005 um^3^. Interestingly, we observed significant differences in the distribution of dots, which suggests large structural heterogeneity of the chromatin at this scale.

**Figure 2.**
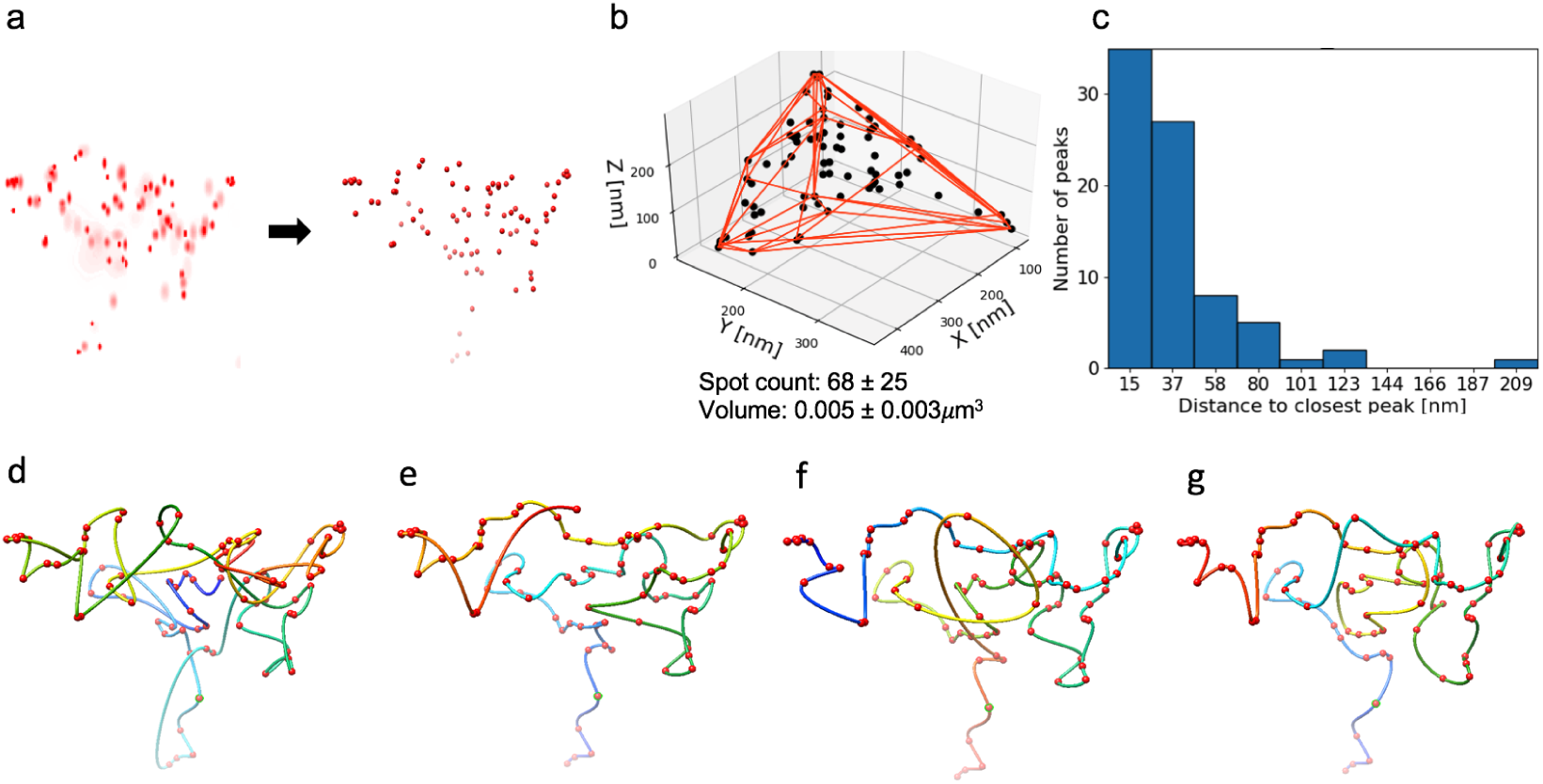
iPALM image-driven chromatin loop models for image 2. a. Dots are identified using iPALM signal processing algorithm in image 2 ***Figure 1***e. The left is fluorescent signals from pre-processed image and the right is identified dots after processing. b. Identified dots with a convex hull represented by red lines are used to calculate the volume estimation for image 2. Dot counts and volume statistics for all six images are calculated. c. Distance histogram between two closest dots in image 2. iPALM image-based chromatin models are produced by four different connecting algorithms. d. neighbor joining, e. nearest neighbor 1, f. nearest neighbor 2, g. traveling salesman. **Figure 2–video 1**. Three-dimensional projection of NJ model for image 1. **Figure 2–video 2**. Three-dimensional projection of NN1 model for image 1. **Figure 2–video 3**. Three-dimensional projection of NN2 model for image 1. **Figure 2–video 4**. Three-dimensional projection of TSP model for image 1. **Figure 2–video 5**. Three-dimensional projection of NJ model for image 2. **Figure 2–video 6**. Three-dimensional projection of NN1 model for image 2. **Figure 2–video 7**. Three-dimensional projection of NN2 model for image 2. **Figure 2–video 8**. Three-dimensional projection of TSP model for image 2. **Figure 2–video 9**. Three-dimensional projection of NJ model for image 3. **Figure 2–video 10**. Three-dimensional projection of NN1 model for image 3. **Figure 2–video 11**. Three-dimensional projection of NN2 model for image 3. **Figure 2–video 12**. Three-dimensional projection of TSP model for image 3. **Figure 2–video 13**. Three-dimensional projection of NJ model for image 4. **Figure 2–video 14**. Three-dimensional projection of NN1 model for image 4. **Figure 2–video 15**. Three-dimensional projection of NN2 model for image 4. **Figure 2–video 16**. Three-dimensional projection of TSP model for image 4. **Figure 2–video 17**. Three-dimensional projection of NJ model for image 5. **Figure 2–video 18**. Three-dimensional projection of NN1 model for image 5. **Figure 2–video 19**. Three-dimensional projection of NN2 model for image 5. **Figure 2–video 20**. Three-dimensional projection of TSP model for image 5. **Figure 2–video 21**. Three-dimensional projection of NJ model for image 6. **Figure 2–video 22**. Three-dimensional projection of NN1 model for image 6. **Figure 2–video 23**. Three-dimensional projection of NN2 model for image 6. **Figure 2–video 24**. Three-dimensional projection of TSP model for image 6.

### Reconstruction of the chromatin target by iPALM image modeling

To better understand the visualized structures, we reconstructed the single chromatin loop from the pre-processed images. We demonstrated a new image processing algorithm ***Figure 2–Figure Supplement 2*** that identifies the coordinates of the single molecules from pre-processed iPALM images (***Figure 1***e) and returns points localizations in a PDB file format (***Figure 2***a), which can be used in further modeling. Basically, we extracted significant signals from the images. We measured the brightness of signals in relative luminosity units which range from 0 to 255. Brightness threshold was chosen manually to cut off noise. In this way, we got different sets of dots for each image.

Number of points varied between images as well, ranging from 42 to 110 which is much smaller than the total number of probe oligos for each chromatin target. Several reasons could cause this under-stained effect: the chromatin is not ideally fully stained due to the FISH efficiency; some signals are lost during data processing.

We used three different algorithms to simulate and reconstruct the chromatin conformation. Each set of dots was connected using Neighbor Joining algorithm (NJ), Nearest Neighbor algorithm with two different starting positions (NN1 and NN2), and Traveling Salesman Problem solver (TSP). The obtained structures were smoothed using cubic spline interpolation (***Figure 2***d,e,f,g). Therefore, we got four probable structure models for each image. We then measured the linear length of these modeled structures (***Table 2***). Considering the probing density is in average of 10 fluorophore/kb, we were able to resolve the “beads on string” chromatin structure with around 150 bp per unit. As previously reported, the first level of DNA compaction from base pair backbone to histone modified “beads on string” structure is about five to ten fold condensation in size (***Felsenfeld and Groudine, 2003***). Therefore, our 33kb target chromatin is estimated to be around 2244 nm to 4488 nm with the first level “beads on string” structure. Compared with our modeled DNA length ***Table 2***, most of them are within the estimated range, only two of the images have slightly shorter length.

**Table 2.** Polymer length in nm

| Method | image 1 | image 2 | image 3 | image 4 | image 5 | image 6 |
| --- | --- | --- | --- | --- | --- | --- |
| NJ | 2569.35 | 3481.72 | 1897.23 | 3218.16 | 1967.29 | 5346.97 |
| NN1 | 2320.08 | 2651.38 | 1562.75 | 3092.21 | 1743.19 | 3829.22 |
| NN2 | 2119.16 | 2730.49 | 1606.68 | 2592.97 | 1742.47 | 4013.24 |
| TSP | 2038.64 | 2596.36 | 1383.92 | 2404.38 | 1417.93 | 3623.48 |

### iPALM image evaluation by comparing image models and Hi-C data

To evaluate iPALM image-driven models, we first compared the distance matrix from 3D model with Hi-C contact matrix using 1kb resolution for 34kb length region (23,016,000 – 23,050,000). Figure 3a shows the distance map of the NN1 model using image 2, as a typical loop structure. The 1kb bead-pairwise distance from 3D model varied from 0nm (darkest blue) to 402nm (dimmest blue or white). Using the Hi-C map as a control (***Figure 3***b), the comparison map was produced in ***Figure 3***c, measuring the fold change of the distance map to Hi-C contact map. To calculate the comparison map, we rescaled the distance map such that the minimum distance is the maximum value while the maximum distance is the minimum value with a linear interpolation, and then multiplying the distance map by the weighting factor so that the average value of the rescaled distance map is the same as the average value in Hi-C contact map across 34 x 34 matrix. White color in the comparison map represent the regions where both maps are similar, and red color those where the distance values are larger than Hi-C contact map values, whereas blue color the regions where Hi-C contact map values are larger than distance value in 3D model. The comparison map shows that the loop region of 3D model is similar to that in Hi-C, while 3D model shows higher value in off-diagonal region. Hi-C is a population averaged contact data from millions of cell, and signals randomly scattered in off-diagonal area or outside TAD regions are treated as noise, reduced by balanced normalization. On the other hand, individual iPALM image-driven models are based on each unique iPALM image, and the distance map shows strong off-diagonal value.

**Figure 3.**
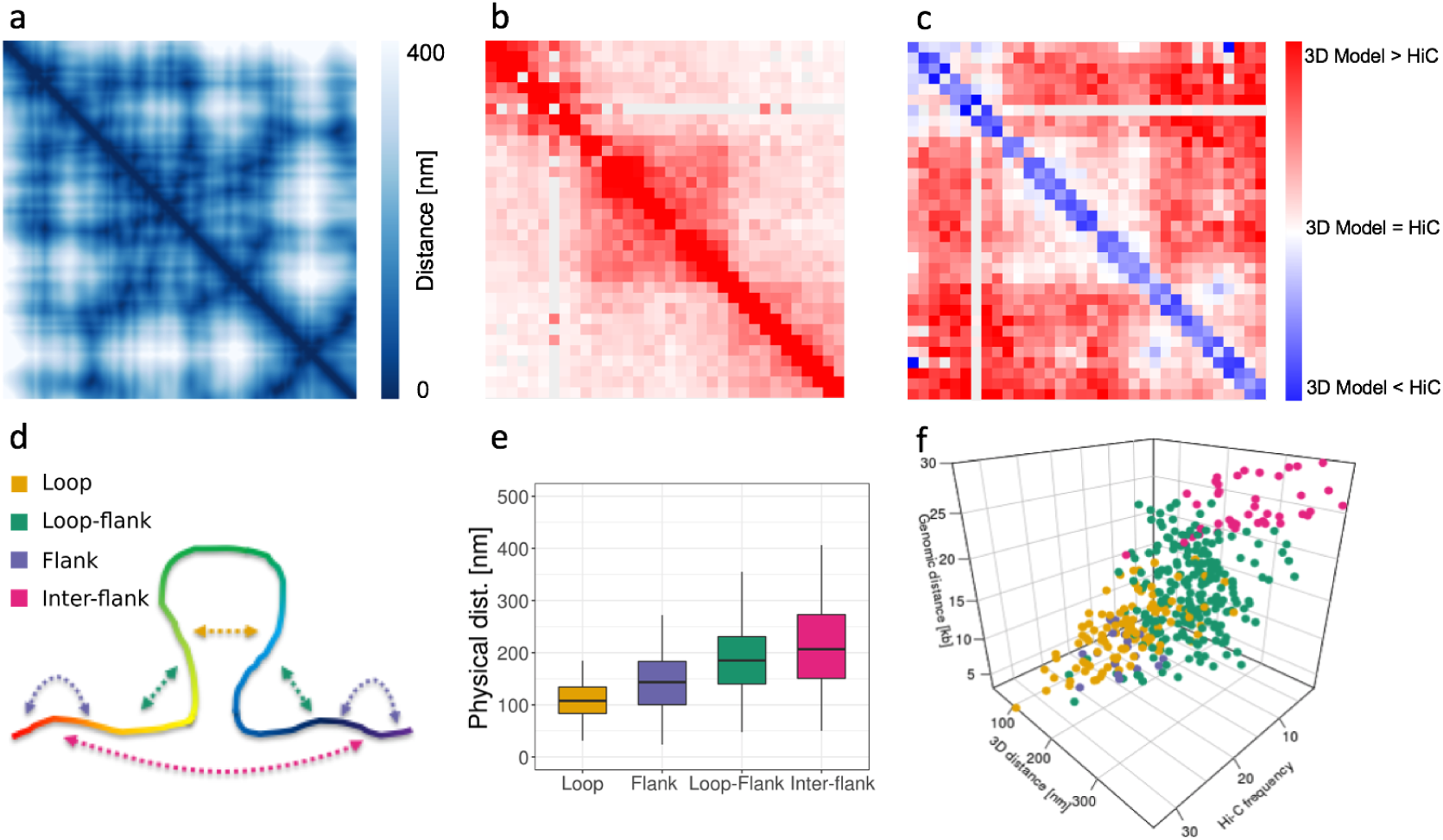
Image-driven model evaluation by comparing with Hi-C data. a. Distance map of NN1 model for image 2 ***Figure 2***e. b. Hi-C contact map (chr14: 23.016-23.05Mb, 1kb resolution). c. Contrast map of ***Figure 3***a. and ***Figure 3***b. d. Illustration of pairwise distance calculation, presenting four interaction groups: intra-loop interaction (yellow), intra-flank interaction (purple), loop-flank interaction (green), inter-flank interaction (red). e. Boxplots for physical distance of four interaction groups for image-driven model in ***Figure 2***e. f. 3D scatter plot presenting the chromatin contact distribution with axis of genomic distance, physical distance, and Hi-C frequency.

From Hi-C, and CTCF and RNAPII ChIA-PET data, we expect the strong 13kb loop region in the middle of target region and 10kb flanking region at each side of the target region. ***Figure 3***d illustrates four different interaction groups in the target region using dotted arrows: intra-loop (yellow), intra-flank (purple), loop-flank (green), and inter-flank (red) for the 1kb bead pairwise physical distance in 3D models. The 3D physical distance from the image-driven model matched well with the genomic expectation, in ***Figure 3***e. The intra-loop and intra-flank distances were significantly lower than those for the loop-flank and inter-flank distances (***Figure 3***e). This effect was preserved when we removed the shortest contacts from the analysis.

For more comprehensive analysis, 3D scatter plot was produced by Hi-C frequency, genomic distance [kb], and physical distance [nm] in ***Figure 3***f. Cleary, it shows that the intra-loop pairs are highest Hi-C frequency and closest physical distance, while pairwise interaction points spread toward lower Hi-C frequency and farther physical distance for intra-flank, loop-flank, and inter-flank, respectively. We generated four different 3D image-driven models for all six iPALM images and showed the results in the supplementary figure 4 – 9 ***Figure 3–Figure Supplement 1, Figure 3–Figure Supplement 2, Figure 3–Figure Supplement 3, Figure 3–Figure Supplement 4, Figure 3–Figure Supplement 5, Figure 3–Figure Supplement 6***. The iPALM image-driven models show the dynamic and heterogeneous chromatin structures, but many models capture the major looping structure as the highest Hi-C frequency and the closest physical distance in the intra-loop interaction group, shown in the supplementary figure (***Figure 3–Figure Supplement 1, Figure 3–Figure Supplement 2, Figure 3–Figure Supplement 3, Figure 3–Figure Supplement 4, Figure 3–Figure Supplement 5, Figure 3–Figure Supplement 6***).

## Discussion

This is the first time a candidate chromatin loop is investigated and visualized using super-resolution microscopy. Though we expected to see a major looping structure in the target chromatin region inferred by ChIA-PET and Hi-C sequencing data, we observed more complex looping structures in each individual image that differentiate them from each other. There is a variety of possible reasons for the observation. 1. The chromatin loop could be heterogeneous between cells and alleles, or more dynamic than static in different cell stages. The captured loop suggested by ChIA-PET data indicates one of the preferred conformations that are occurring most frequently at that region. It does not mean that other conformations could not happen as those could be too rare to be captured. Our imaging data is not inclusive enough due to the limitation of the sample size to reflect the frequencies of each type of the looping incidences. 2. There are structures that cannot be captured by molecular strategies. Both ChIA-PET and Hi-C are based on a hypothesis that the chromatin interactions are mediated by protein factors. In other words, the chromatin structure that has no protein binding cannot be captured and modeled, but we cannot deny there are ultra complex twisting and tangling for DNA packing in the nucleus. This study allows a direct visualization of the chromatin looping in a specific region that we clearly see the physical winding of DNA, though there are limitations hindering us from interpreting the observations more comprehensively. For instances, we are not able to exactly link the iPALM images to the corresponding genome coordinates; we could not image the non-looping regions inferred by genome sequencing data; a larger sample size would be helpful bridging the gap of the comparison between individuals by imaging and populations by sequencing.

The imaging data is more direct and straightforward for revealing chromatin conformation than the sequencing data. Therefore, the observations or findings that are against or not in the agreement with our assumptions from the sequencing data are not unexpected, but even more suggestive to the current understanding the chromatin looping.

## Methods and Materials

### Cell culture

GM12878 cells were cultured in RPMI 1640 with 2mM L-glutamine and 15% fetal bovine serum at 37 °C.

### Nuclei isolation

Nuclei EZ Prep Nuclei Isolation Kit from Sigma was used to isolate nuclei from GM12878 cells.

### Coverslip coating

Coverslips were incubated in 1M KOH for 20min, washed with water, coated with 0.01% poly-L-lysine (Sigma) for 20min, rinsed with water, dried for 30min for later use.

### DNA FISH

Isolated nuclei were added to attach to poly-L-lysine coated coverslips, fixed with 4% PFA for 10min at room temperature, washed with 1xPBS, permeabilized with ice-cold methanol for 10min, washed with 1xPBS, dehydrated with 75%, 85% and 100% ethanol for 2min each, dried at 60°C for 1h. FISH probes were mixed with hybridization buffer and added to prepared nuclei, denatured at 80°C for 5min, incubated in a humid chamber at 37°C for overnight. Nuclei were washed with 50% formamide/2xSSC at room temperature for 10min, followed by 2xSSC for 10min, 0.2xSSC at 55°C for 10min and then large volume of 2xSSC till imaging.

### iPALM imaging

Samples were imaged in standard stochastic optical reconstruction microscopy (STORM) buffer by iPALM (***Shtengel et al., 2009***). Isolated nuclei were adhered to 25mm round coverslips containing gold nanorod particles that act as calibration standards and alignment/drift fiducial markers. These were prepared as described in ***Shtengel et al***. (***2014***). Briefly, coverslips were washed for 3 hours at 80 degrees C in a 5:1:1 solution of H2O:H2O2:NH3OH, rinsed copiously, and coated with poly-L-lysine. After further washing, gold nanorods (Nanopartz, Inc) were adhered to poly-L-lysine coated coverslips, washed again, and coated with ca. 50nm SiO2 using a Denton vacuum evaporator. Samples were mounted in dSTORM buffer (***Dempsey et al., 2011***), containing tris buffered saline, pH 8, 100mM mercapto ethanolamine, 0.5 mg/mL glucose oxidase, 40 ug/mL catalase, and 10% (w/v) glucose (all from Sigma). An 18mm coverslip was adhered atop the bottom coverslip, sealed, mounted in the iPALM, and imaged as described above.

### Image pre-processing

iPALM images were reconstructed via localization of blinking fluorophores over 25,000 frames across each of three EMCCD cameras. Gold nanoparticles act as fiducial markers that allow for (1) spatial registration of the three cameras using a full affine transformation, and (2) calibration of the z-position response of the system. After localization, images were filtered to only include localizations with <30nm uncertainty in all three dimensions. The gold fiducial particles also allow for drift correction in all three dimensions, and for correcting any sample tilt to within 30nm error. Final images were rendered as image stacks, reflecting the fluorophore density and uncertainty of localization, or exported as ASCII delimited text files for further analysis.

### Dots identification algorithm

Here we propose the dot identification algorithm implemented to analyze post processed iPALM images. It analyzes three dimensional TIFF files in order to find coordinates of all visible dots (probe oligos attached to the chromatin). Based on a manually set brightness threshold it builds a three dimensional graph, represented as three dimensional matrix, where voxels of image are nodes. Edges are created between two voxels that are located next to each other and their brightness is higher than given threshold. Then the algorithm identifies all connected components in such graph. We define connected component as a part of the image in which all voxels form an area with brightness above cutoff level, which means that they are represented by nodes connected by vertices in created graph. In each connected component we identify a point by finding the XYZ coordinates of the brightest voxel. List of identified dots is remembered, brightness level is increased by a step size, new graph is created and the whole procedure is repeated until the algorithm reaches given maximal brightness level ***Figure 2–Figure Supplement 2***. When the maximal brightness level is reached all identified dots are merged into one set (to avoid repetitions) and list of coordinates of identified dots is returned and saved in PDB format. To the best of our knowledge this is the first approach to predict the dot location from 3D TIFF images.

### Dot-joining algorithms

To comprehensively model the potential chromatin loop structure, we applied three different algorithms to render the models. After that we applied spline interpolation to smooth the models.

#### Neighbor Joining algorithm

Neighbor Joining is an agglomerative clustering method used in bioinformatics for creation of phylogenetic trees. In our approach in each step the algorithm is searching for a pair of dots that are the closest to each other and connecting them in one. This step is repeated until there is no unconnected dots left. We used this approach to connect sets of identified dots from iPALM images.

#### Nearest Neighbor algorithm

Nearest Neighbor algorithm is solving shortest path problem. Shortest path problem in graph theory is the problem of finding a path between two nodes such that the distance between them is minimized. Algorithm is starting from given dot, searching the nearest dot in surroundings, connecting them and again searching for the nearest neighbor and this way connecting a whole set of dots. After analyzing the genomic data we found out that examined loop is in between two subdomains. We assumed that flanking regions will be far apart from each other with the loop in the middle. We run two NN simulations each starting from one of two dots that were the furthest apart in space. Thus, this approach will generate two possible image models.

#### Travelling Salesman algorithm

We used implementation of greedy algorithm finding one of the best solutions for this NP-hard problem. We treat our set of points as graph nodes, and distances between them as vertices. At the beginning each vertex is a separate path of length 1. In each step we are finding two closest disconnected paths, and we connect them into one. This step is repeated until there is just one path left. Greedy TSP solver gives highly non optimal results therefore after connecting all paths into one we run optimization algorithm. Optimization tries to rearrange dots in the path to improve the solution. After finding the shortest path we are simply deleting the longest connection between two dots. This way we get the shortest path between two dots that are the furthest from each other.

### Spline Interpolation

Spline interpolation is class of interpolation which uses polynomials to create smooth function on interval [*a, b*]. This interval is split into m sub-intervals such as *a* = *t*_0_ *< t*_*i*_ *<* … *< t*_*m*_ = *b*. For each point *t*_*i*_ there is a defined value *y*_*i*_ throughout that the interpolation should go. For each of these intervals a different polynomial is used as defined, so they are connecting to a continuous function. The spline degree d uses such polynomials degree at maximum of d to satisfy the condition that derivatives on the whole interval [*a, b*] up to level *d* −1 are continuous. In our case we decided to use cubic splines. The degree limit is set to three.

We set a polynomial *P*_*i*_ on interval [*t*_*i*_, *t*_*i*+1_], and got the following conditions:

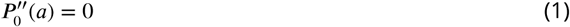

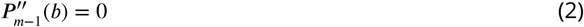

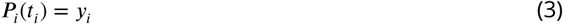

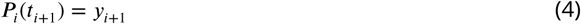

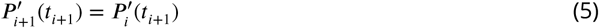

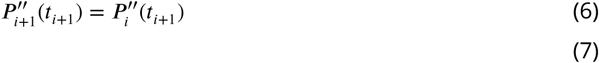

We chose cubic spline interpolation because it gives an interpolating polynomial that is smoother and easier to compute than other methods. We used it to smooth structures modeled from iPALM images.

### Multidimensional scaling

Multidimensional scaling (MDS) algorithm is a statistical method which takes matrix of similarities or distances between objects and put that objects in N-dimensional space possibly close to given distances (***Borg et al., 2017***). In particular we can use Hi-C/ChIA-PET relative frequency contact matrix into physical distances and seek for its representation in 3D space. In this case every bin in interaction matrix will represent a single bead in of a model in 3D. We used Hi-C contact matrix for studied region to obtain 3D chromatin models from genomic data. This matrix we interpreted as graph neighborhood matrix. Using this graph we calculated graph distance. The result was an input to MDS algorithm (Scikit implementation, ***Pedregosa et al***. (***2011***)).

## Acknowledgements

iPALM imaging was done in collaboration with the Advanced Imaging Center at Janelia Research Campus, a facility jointly supported by the Gordon and Betty Moore Foundation and the Howard Hughes Medical Institute. This work was carried out within the TEAM program of the Foundation for Polish Science (TEAM to D.P.) co-financed by the European Union under the European Regional Development Found, co-supported by OPUS grant from Polish National Science Centre (2014/15/B/ST6/05082) and the grant 1U54DK107967-01 “Nucleome Positioning System for Spatiotemporal Genome Organization and Regulation” within 4D Nucleome NIH program. The work was co-supported by the European Commission as European Cooperation in Science and Technology COST actions: CA18127 “International Nucleome Consortium” (INC), and CA16212 “Impact of Nuclear Domains On Gene Expression and Plant Traits”. PS was supported by a PhD stipend from Polish National Science Center (2016/20/T/NZ2/00511).

---

**Figure 1–Figure supplement 1.**
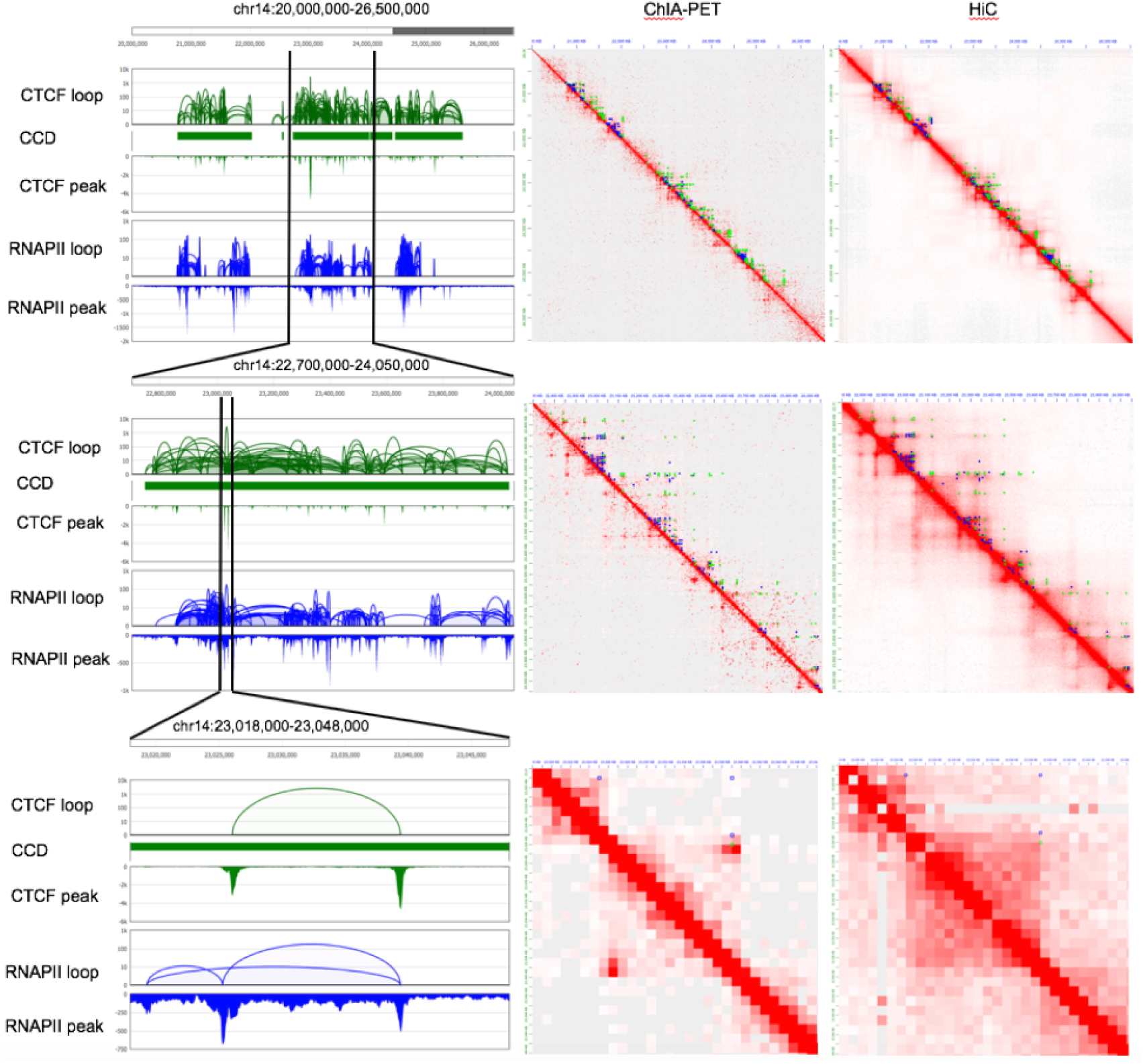
Multi-scale view of the selected target loop region in GM12878. Three different scales over the target loop region are presented from the zoomed out view of 6.5Mb length (top row, Chr14: 20 – 26.5Mb), 1.35Mb length (middle row, Chr14: 22.7 – 24.05Mb), to the zoomed in view of 30Kb length (bottom row, Chr14: 23.018 – 23.048Mb). The first column is the genome browser view of CTCF and RNAPII ChIA-PET loops and peaks with CCDs (CTCF-mediated chromatin contact domains). The second and third column are 2D contact maps of ChIA-PET (CTCF and RNAPII merged) and Hi-C, respectively. ChIA-PET loops (green for CTCF and blue for RNAPII) are also presented on top of the upper diagonal area of the both ChIA-PET and Hi-C contact maps. Balanced normalization was applied to contact maps, and three different resolutions were used, 10kb (top row), 5kb (middle row), and 1kb (bottom row).

**Figure 2–Figure supplement 1.**
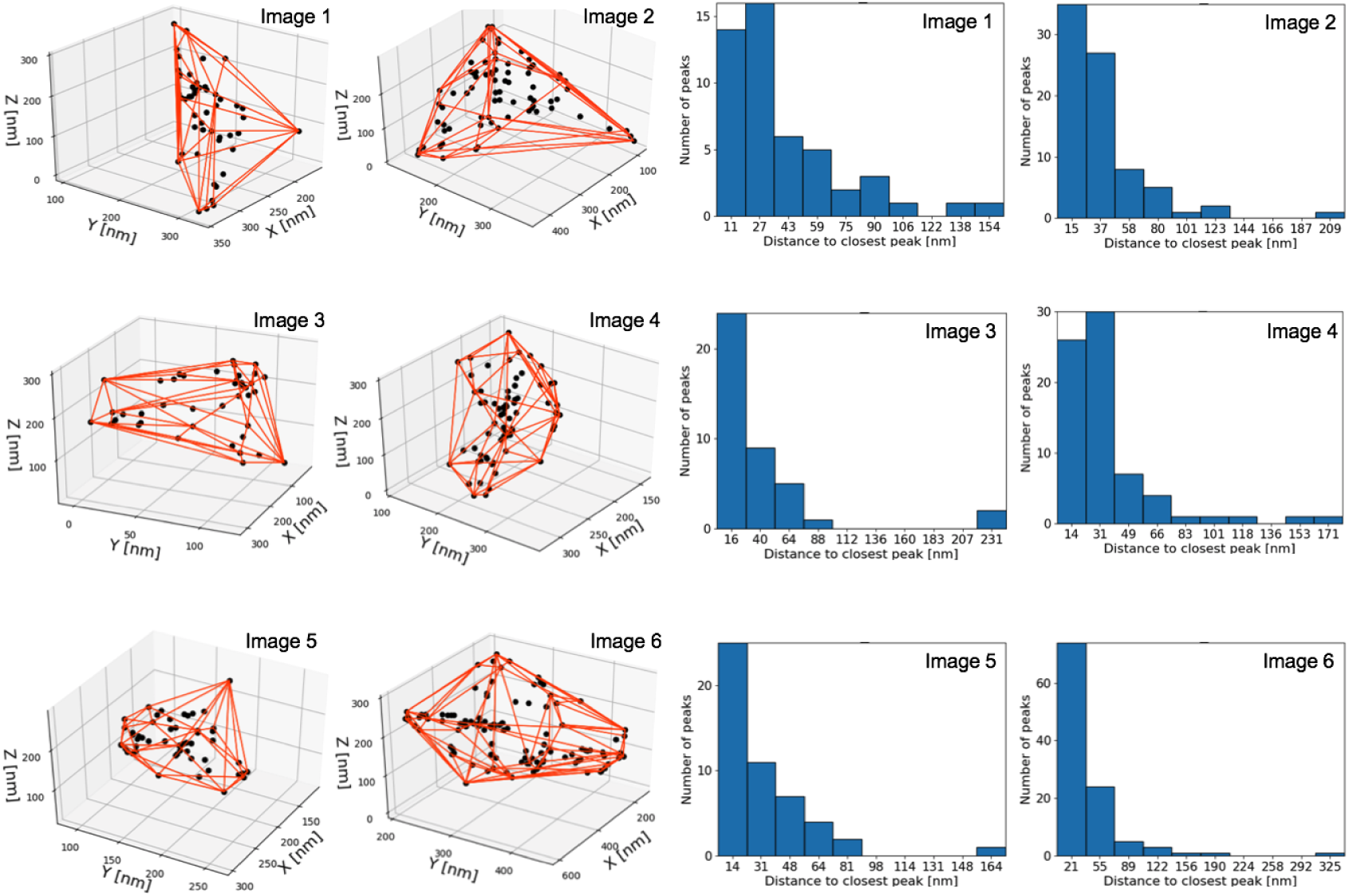
Identified iPALM image dots for all six images. Identified dots with convex hull (red lines) and distance histogram between two closest dots for all six images.

**Figure 2–Figure supplement 2.**
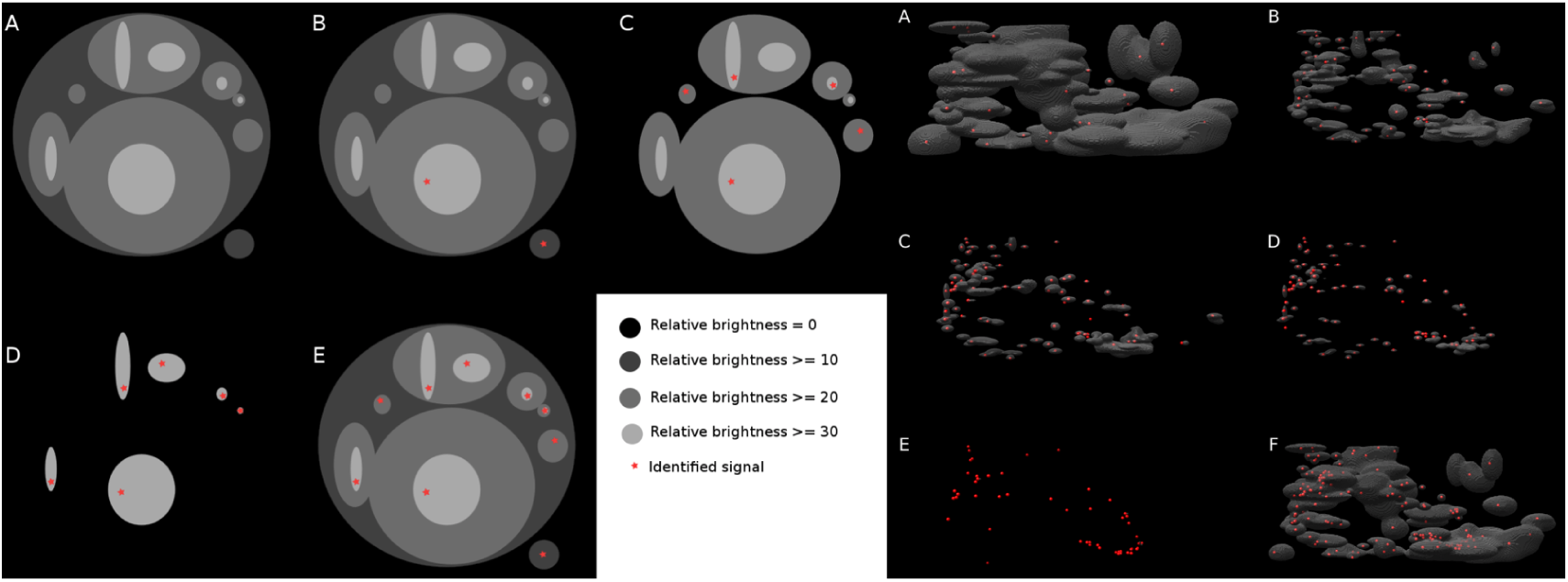
iPALM signal processing algorithm. The left panel is a schematic illustration to extract dots from the broad signal density image. Algorithm starts from the lowest brightness level and identifies connected components (A) in each connected component the brightest voxel is identified (B) then the brightness level is changed by a step size and the connected component analysis and brightest voxel identification is repeated until reaching the maximal brightness of the image (C,D). At the end all of identified voxels are connected into one set to avoid repetitions. Intermediate processed screenshots for image 6 were shown in the right panel. Brightest voxels identification in connected components found at different brightness levels (A,B,C,D,E) and all identified voxels merged into one set (F).

**Figure 3–Figure supplement 1.**
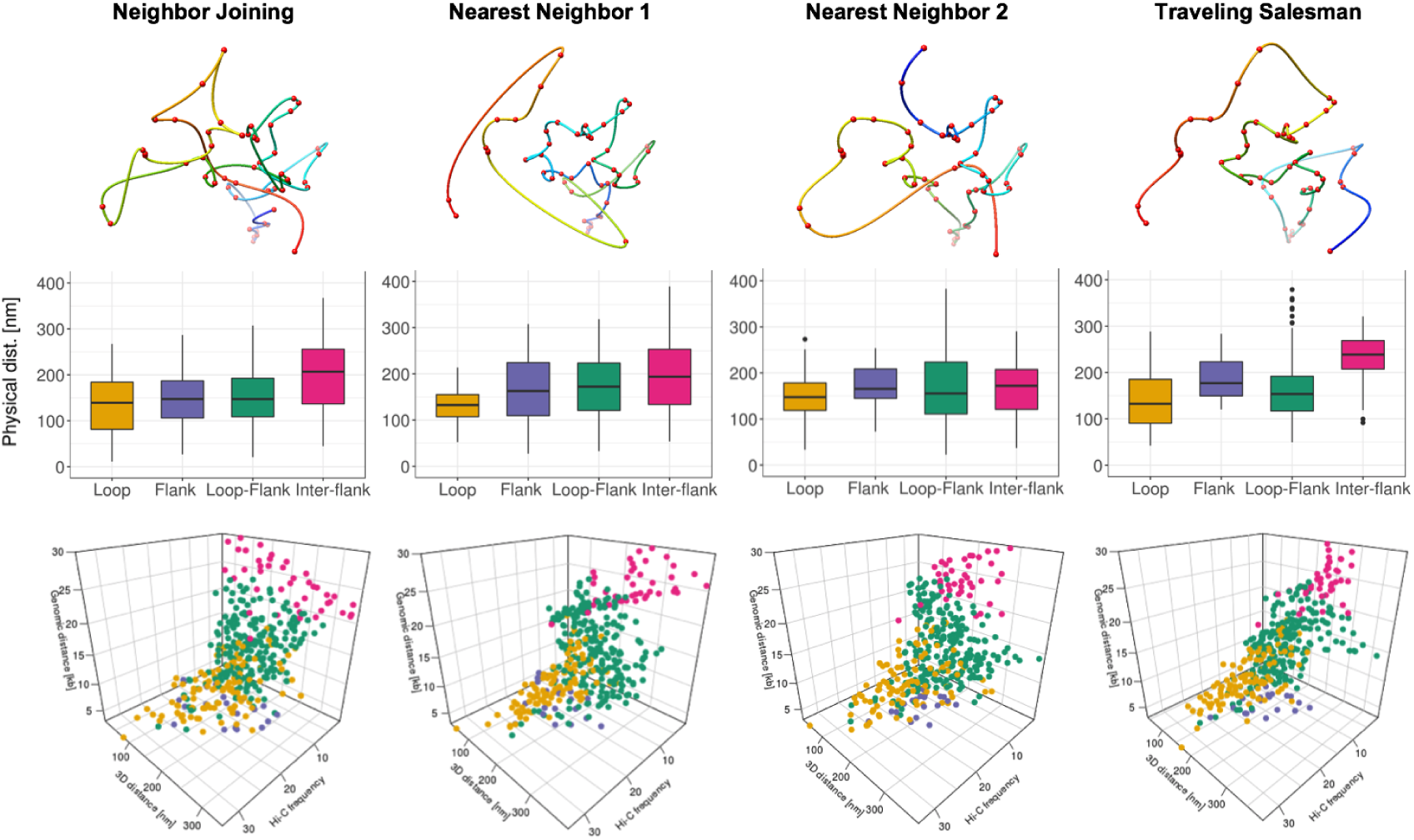
Image-driven 3D models, boxplots, and 3D scatter plots for four interaction groups (image 1).

**Figure 3–Figure supplement 2.**
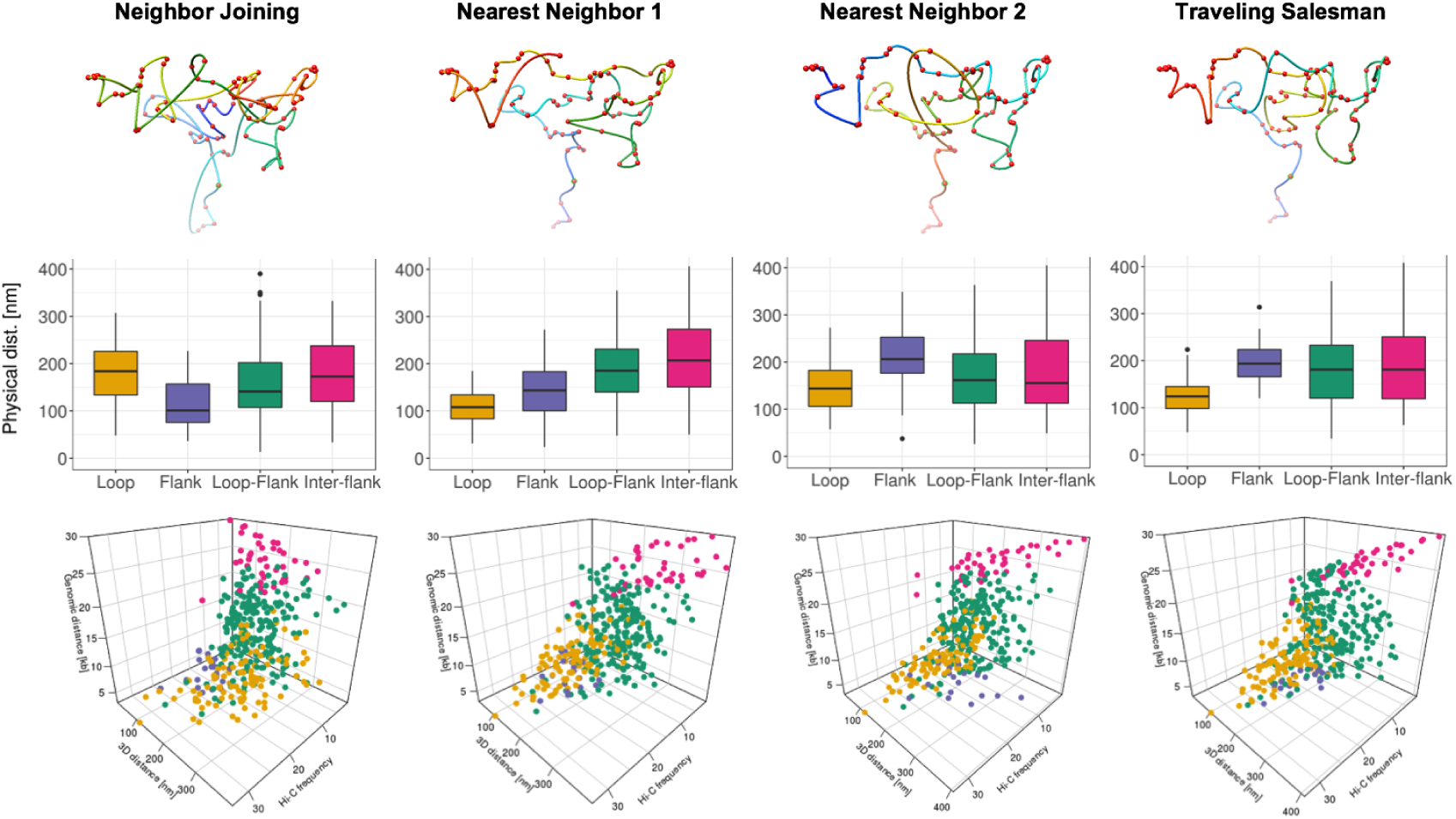
Image-driven 3D models, boxplots, and 3D scatter plots for four interaction groups (image 2).

**Figure 3–Figure supplement 3.**
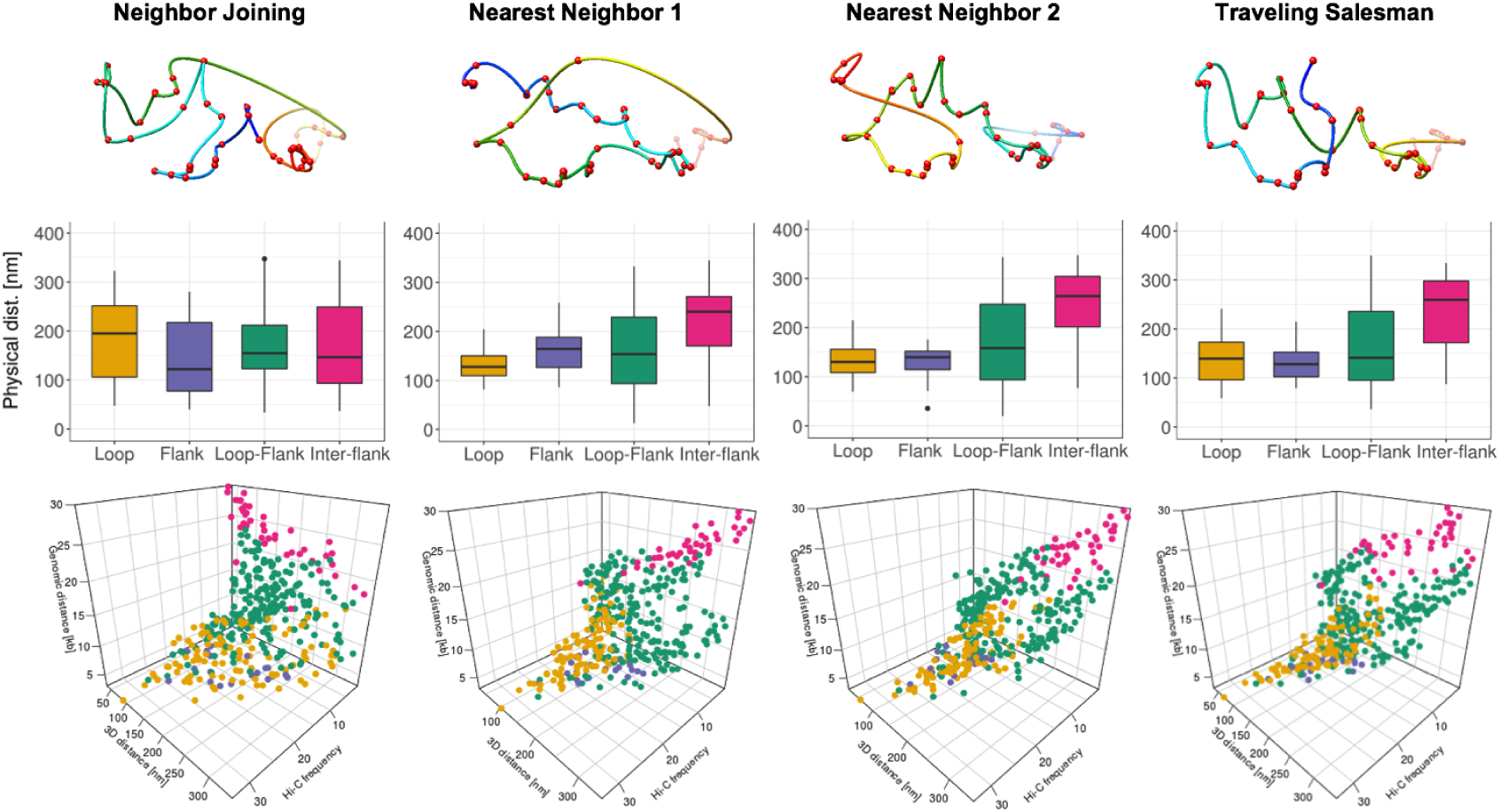
Image-driven 3D models, boxplots, and 3D scatter plots for four interaction groups (image 3).

**Figure 3–Figure supplement 4.**
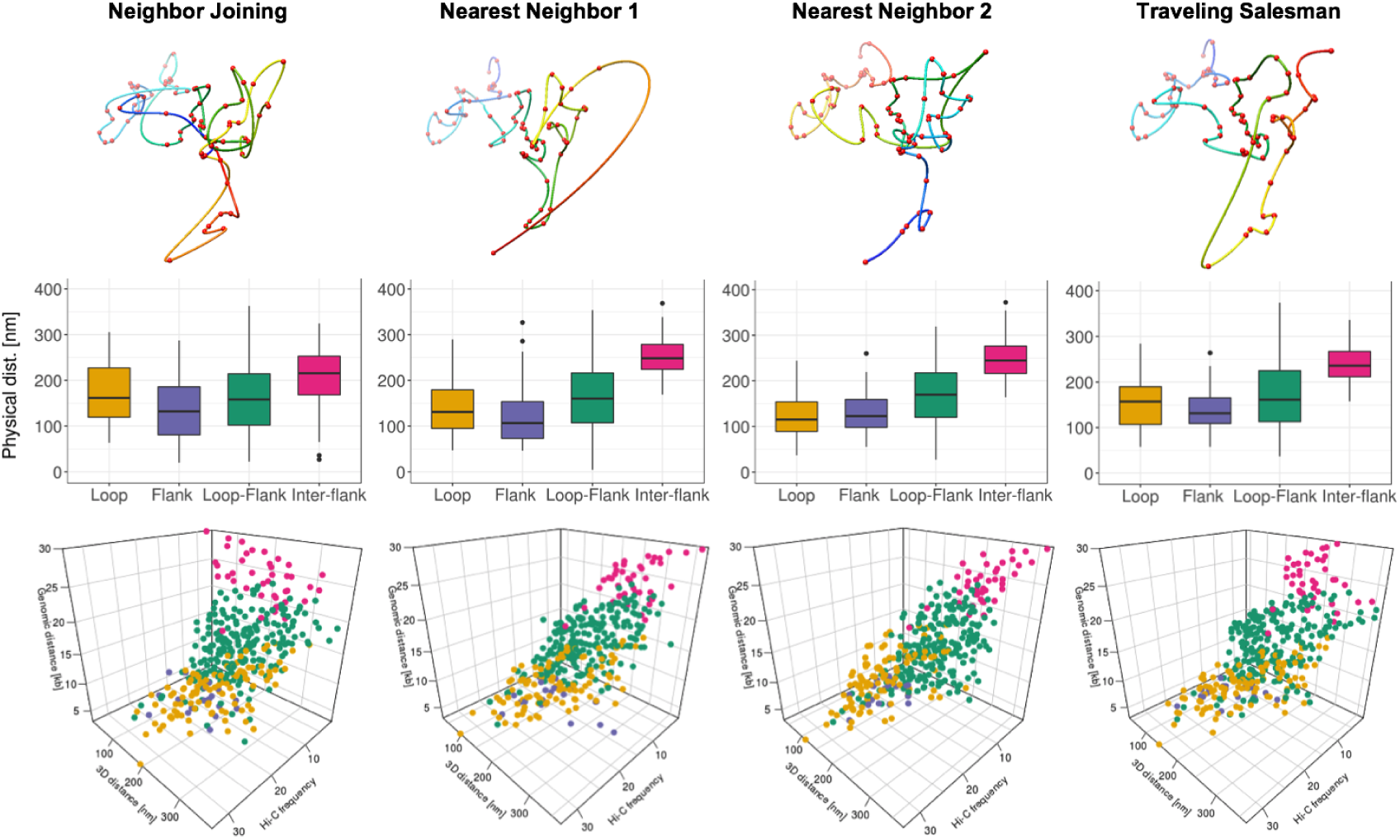
Image-driven 3D models, boxplots, and 3D scatter plots for four interaction groups (image 4).

**Figure 3–Figure supplement 5.**
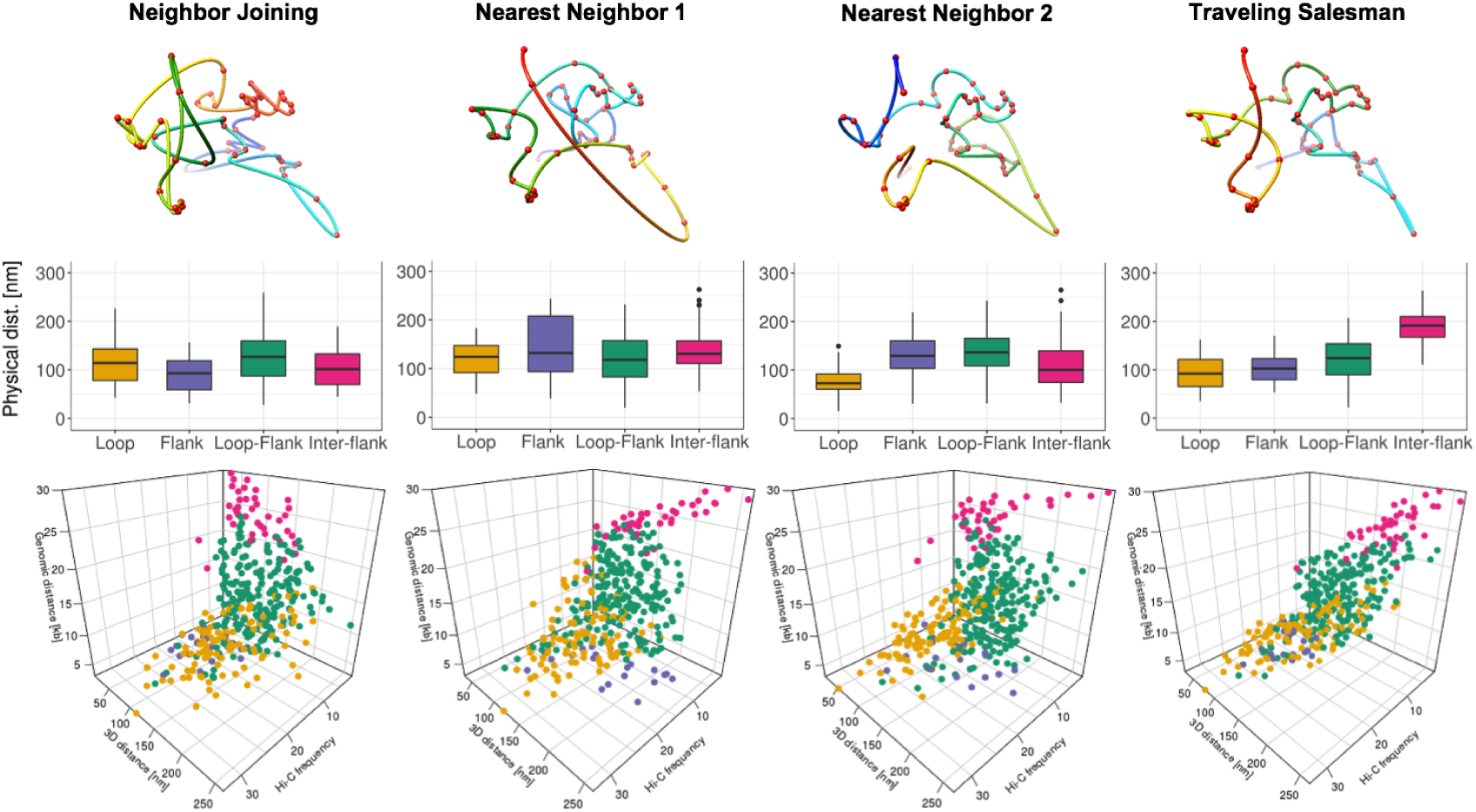
Image-driven 3D models, boxplots, and 3D scatter plots for four interaction groups (image 5)

**Figure 3–Figure supplement 6.**
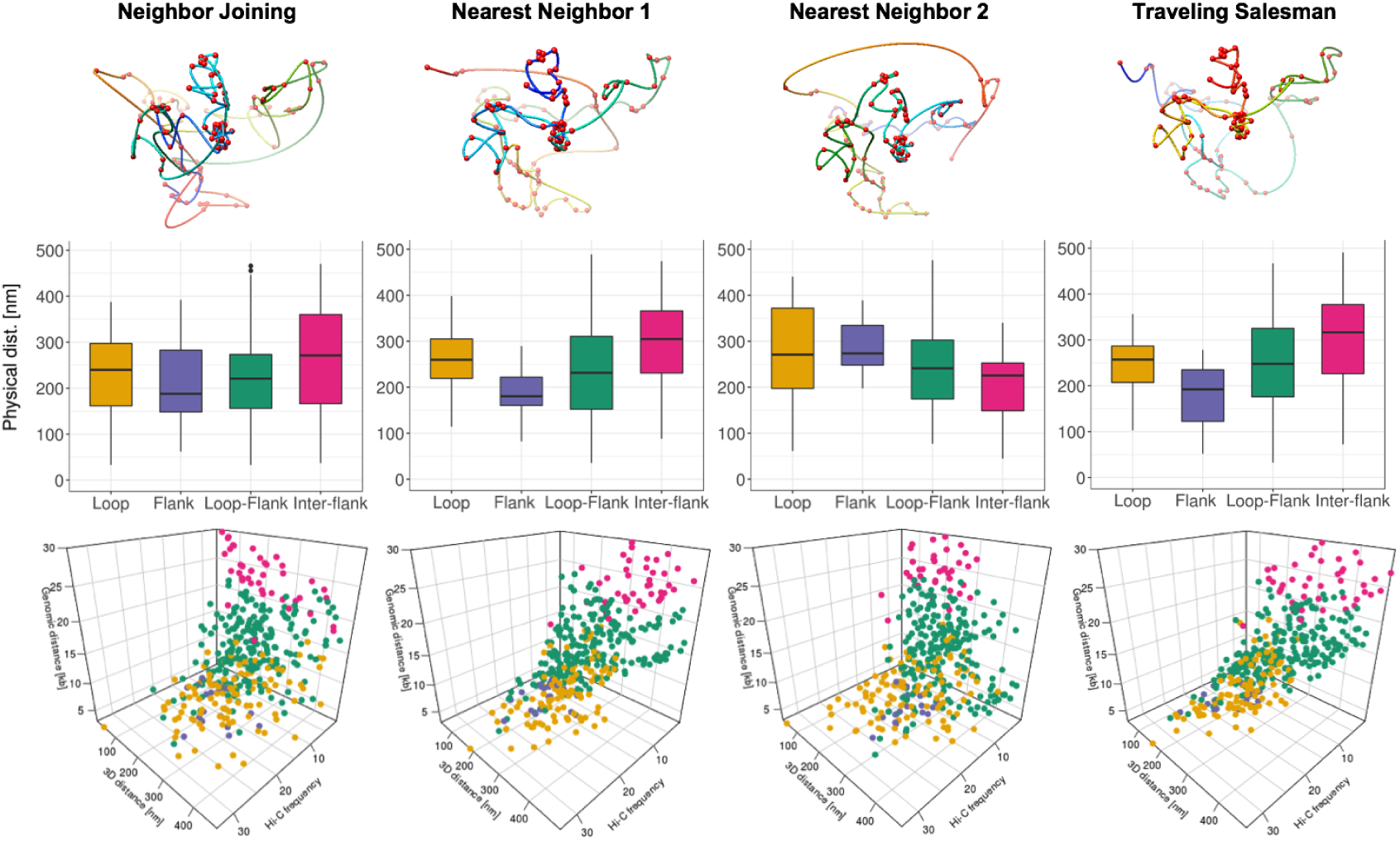
Image-driven 3D models, boxplots, and 3D scatter plots for four interaction groups (image 6).

